# Bimodal Specificity of TF-DNA Recognition in Embryonic Stem Cells

**DOI:** 10.1101/2024.09.18.613654

**Authors:** Michael Povolotskii, Maor Yehezkehely, Oren Ram, David B. Lukatsky

## Abstract

Transcription factors (TFs) bind genomic DNA regulating gene expression and developmental programs in embryonic stem cells (ESCs). Even though comprehensive genome-wide molecular maps for TF-DNA binding are experimentally available for key pluripotency-associated TFs, the understanding of molecular design principles responsible for TF-DNA recognition remains incomplete. Here, we show that binding preferences of key pluripotency TFs, such as Pou5f1 (Oct4), Smad1, Otx2, Srf, and Nanog, exhibit bimodality in the local GC-content distribution. Sequence-dependent binding specificity of these TFs is distributed across three major contributions. First, local GC-content is dominant in high-GC-content regions. Second, recognition of specific *k*-mers is predominant in low-GC-content regions. Third, short tandem repeats (STRs) are highly predictive in both low- and high-GC-content regions. In sharp contrast, the binding preferences of c-Myc are exclusively dominated by local GC-content and STRs in high-GC-content genomic regions. We demonstrate that the transition in the TF-DNA binding landscape upon ESC differentiation is solely regulated by the concentration of c-Myc, which forms a bivalent c-Myc-Max heterotetramer upon promoter binding, competing with key pluripotency factors such as Smad1. Finally, a direct interaction between c-Myc and key pluripotency factors is not required to achieve this transition.

**Graphical abstract:** 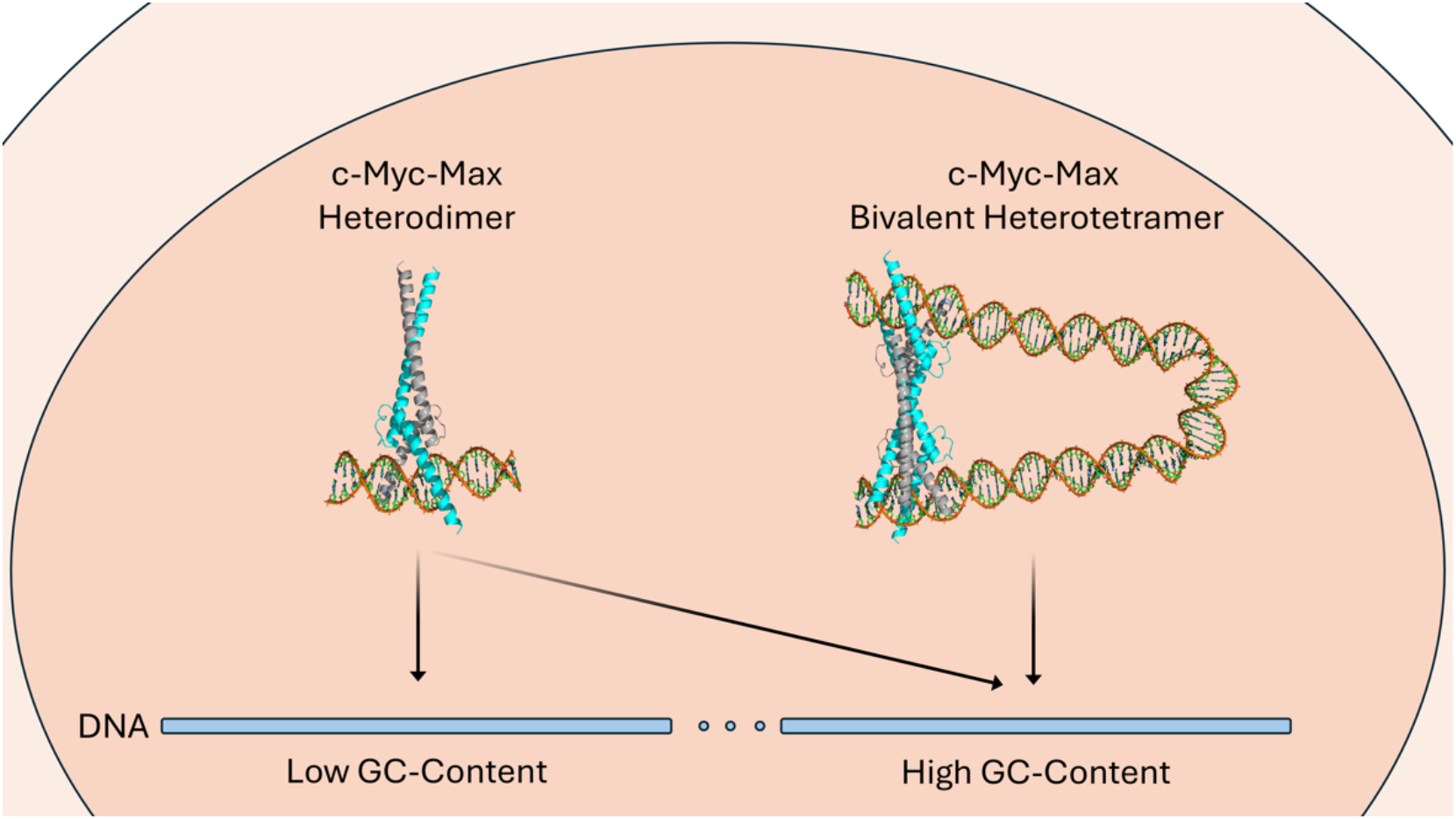

## Introduction

Transcription factors (TFs) bind genomic DNA regulating gene expression (1-7). Using *E*.*coli lac* repressor and λ phage repressor as model systems, seminal studies by Jacob-Monod (2) and Ptashne (7) established that short regulatory DNA sequences specifically bind TFs with high affinity. The subsequent studies by Riggs (4) and von Hippel (5) revealed that in the case of *lac* repressor, 98% or more of repressor is bound to DNA predominantly at sites other than the *lac* operator. It is explained by the fact that inducers shift repressor from operator to nonoperator DNA for significant distances, but do not free it from DNA.

Currently, five decades after those studies, high-throughput methods, such as chromatin immunoprecipitation followed by massively parallel sequencing (ChIP-seq) allow genome-wide identification of TF-DNA binding maps (8-13). Similar to how *lac* repressor binds nonoperator DNA, these maps reveal that the majority of variable TF-DNA binding events occur outside of known specific TF bindings motifs (8,9).

Even though molecular mechanisms responsible for specific TF-DNA recognition have been studied over the past sixty years, many questions still remain unanswered (14-16). Understanding such mechanisms in eukaryotic cells is further complicated by the effect of chromatin structure, promoter-enhancer interactions, protein-protein interactions, and epigenetic modifications of histones and DNA (9,17,18).

Interestingly, genome-wide association studies (GWAS) show that the vast majority of disease-associated genetic variants are located in non-coding regions (17). Understanding of molecular mechanisms responsible for TF recognition of such non-coding variable genomic regions constitutes one of the key open questions impeding the understanding of phenotypic consequences of disease-associated variants (17).

It is of particular importance to establish the link between TF-DNA recognition and gene expression programs in embryonic stem cells (ESCs) due to the similarities between ESCs with various cancers (12). Specific TF-DNA binding motifs provide only a limited understanding of the experimentally determined TF-DNA binding network in ESCs (8). Mechanistic understanding of this binding network is further complicated by the fact that different pluripotency TFs interact with each other forming multimeric protein complexes (12). One such TF, c-Myc, is an important transcriptional regulator in ESCs, somatic cell reprogramming, and various cancers (12). In particular, increased c-Myc expression level constitutes a common feature of undifferentiated ESCs and cancer cells (12). In the past, it was revealed that the c-Myc-centered regulatory module is largely uncoupled, both in terms of protein-protein and TF-DNA binding, from the module comprising other key pluripotency factors, such as Nanog, Pou5f1 (Oct4), Sox2, Smad1 (12). In particular, it was shown that these two modules bind different genomic regions (12). Molecular recognition mechanisms responsible for such behavior remain unknown.

Here, using human embryonic stem cells as a model system, we show that key pluripotency TFs possess bimodal intrinsic DNA recognition specificity characterized by fundamentally different DNA binding mechanisms in low- and high-GC-content genomic regions. In particular, in GC-poor genomic regions, TF-DNA binding recognition is predominantly determined by specifically recognized *k*-mers. In contrast, in GC-rich genomic regions TFs recognize the local genomic GC-content but possess a weak *k*-mer specificity. Non-consensus repetitive elements contribute to TF-DNA recognition in both low- and high-GC-content regions.

We also demonstrate that upon developmental transitions, c-Myc can influence the TF-DNA recognition mode of key pluripotency factors, such as Smad1, Pou5f1 (Oct4), Otx2, and Srf. We provide further evidence that c-Myc competes with these pluripotency factors in GC-rich genomic regions, such as the vicinity of transcription start sites (TSSs). Using a simple chemical kinetics model, we predict that no direct interaction between c-Myc and the module comprising key pluripotency factors, such as Smad1, is required in order to achieve the transition in the TF-DNA binding landscape upon ESC differentiation. Rather this transition is solely regulated by the concentration of c-Myc, competing with other key pluripotency factors for promoter binding. We predict that the formation of a bivalent c-Myc-Max heterotetramer, for example, by looping DNA, is necessary for achieving this transition.

## Materials and Methods

### MNChIP-seq data acquisition

We used micrococcal nuclease (MNase) based chromatin immunoprecipitation sequencing (MNChIP-seq) data obtained in undifferentiated human ESCs and during early stages of ESC differentiation: Mesendoderm (dMS), Endoderm (dEN), Mesoderm (dME), and Ectoderm (dEC) (8). The data used in our analyses are publicly available at the Gene Expression Omnibus under the accession number GSE61475 (8). For the genomic analysis we used the human reference genome version hg19.

### AlphaFold 3 modeling of c-Myc-Max-DNA complex

We used AlphaFold 3 (19) for predicting c-Myc-Max-DNA complex structures for a set of genomic DNA sequences (Supplementary Table S1). In simulations we used truncated protein sequences of c-Myc and Max, representing the DNA-binding domain (DBD), analogous to the sequences used in experimentally obtained c-Myc-Max-DNA crystallographic structures (20) (Supplementary Table S1).

## Results

### Dissecting TF-DNA binding specificity in relation to GC-content bimodality

Statistical analysis of the local GC-content of MNChIP-seq peaks shows a large degree of variability within and between different TFs (Figure 1A). Key pluripotency factors, such as Nanog, Pou5f1 (Oct4), Sox2, Otx2, Smad1, and Srf are characterized by a median GC-content below 0.5 in ESCs (Figure 1A). Remarkably, a significant number of TFs show bimodality in the GC-content distribution (Figure 1B, Supplementary Figure S1). For example, Pou5f1, Smad1, Otx2, Srf exhibit a transition from lower to higher GC-content upon developmental transition from ESCs to dEN (Figure 1B, Supplementary Figure S1). Other TFs, such as Nanog, exhibit a static bimodality in the GC-content distribution of MNChIP-seq peaks at different stages of differentiation (Supplementary Figure S1). On the contrary, c-Myc is characterized by a stable local GC-content distribution with the highest median GC-content (Figure 1A and B). Statistically, genomic regions around transcription start sites (TSSs) represent high-GC-content regions, as well as regions with high binding-peak intensity (Figure 1C). For example, the mean peak intensity of c-Myc and Pou5f1 binding peaks aligned around TSSs correlates with the mean local GC-content (Figure 1C).

**Figure 1.**
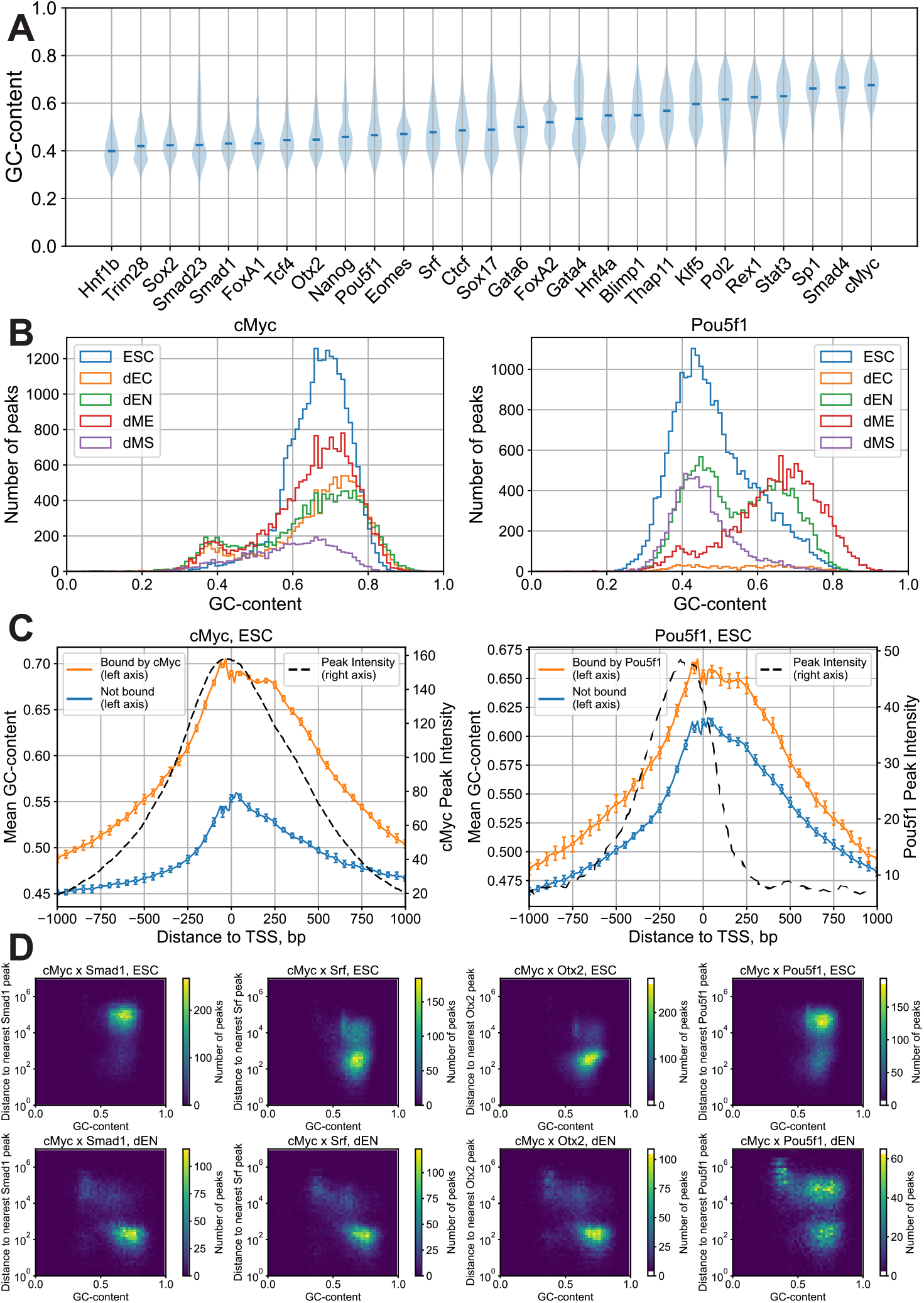
Dissecting TF-DNA binding specificity in relation to GC-content bimodality. (**A**) Violin plot of the GC-content distribution of MNChIP-seq peaks for 27 TFs in ESCs (data is taken from ref. (8)). TFs are sorted by increasing median GC-content of peaks. (**B**) Example: Histogram of the GC-content distribution of MNChIP-seq peaks for c-Myc and Pou5f1 (Oct4) in different developmental stages. (**C**) Correlation between the mean local GC-content and the mean binding intensity of aligned MNChIP-seq peaks for c-Myc and Pou5f1 in ESCs as a function of genomic position relative to TSS. Orange curves show the mean GC-content for TSSs bound by c-Myc (left) and Pou5f1 (right). Blue curves show the mean GC-content for TSSs not bound by c-Myc (left) and Pou5f1 (right). We defined a given TSS as ‘bound’ if it contains at least one binding peak within 1000-bp around the TSS. The local GC-content was computed within a sliding window with a width of 50-bp. For cMyc, the GC-content of bound TSSs shows a strong correlation with the average binding peak intensity, with the Pearson linear correlation coefficient *R* = 0.9*7* (*p* = 1.8 × 10^−118^). For Pou5f1, the correlation is weaker, with *R* = 0.*6*2 (*p* = 1.9 × 10^−22^). Error bars represent one standard deviation of the mean GC-content, determined by dividing the entire set of TSSs into 10 equally sized subgroups and calculating the average GC-content for each subgroup. (**D**) The joint distribution of the GC-content in c-Myc binding peaks and their proximity to the nearest peaks of Smad1, Srf, Otx2, and Pou5f1, respectively, shown for two cell types, ESC and dEN. The distances are measured between peak centers. We partitioned the GC-content range (0 to 1) into 50 equal bins and the distance range (1 to 10^7^-bp) into 49 logarithmically scaled bins, counting the frequency of binding peaks within the bins.

The analysis of co-binding between c-Myc and other key pluripotency factors such as Smad1, Srf, Otx2, and Pou5f1 shows that in the course of differentiation, there is a competition between c-Myc and these TFs for binding GC-rich DNA regions (Figure 1D, Supplementary Figure S2). In particular, upon differentiation from ESCs to dEN, peaks of these TFs undergo a transition towards high-GC-content genomic regions, which were previously occupied by c-Myc (Figure 1D, Supplementary Figure S2). In what follows we show that this transition does not require a direct interaction of c-Myc with these TFs.

### Bimodality of *k*-mer specificity is linked to bimodality of GC-content

The second feature responsible for TF-DNA binding specificity we consider (in addition to the local GC-content) is short genomic sequences of length *k*. We term such sequences *k*-mers. To assess TF specificity to *k*-mers, we developed a peak-over-background statistical energy model, according to which, the binding energy (in the units of *k*_B_*T*, where *k*_B_ is the Boltzmann constant and *T* is the temperature) of a particular *k*-mer named *s* is calculated as follows:

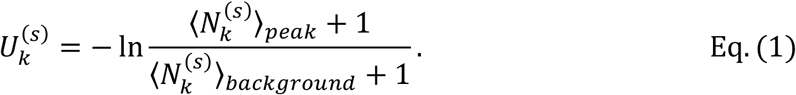

Here,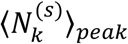 denotes how many times, on average, the *k*-mer *s* and its reverse complement occur in the 100 bp-long nucleotide sequences at the center of MNChIP-seq binding peaks. The denominator, 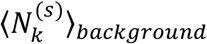 is computed for control sequences, which come from 100-bp-long genomic regions situated 100-bp from the binding peak edges, and not overlapping with other peaks. Adding 1 to both nominator and denominator avoids taking the logarithm of zero and dividing by zero.

For a given TF in a given developmental stage, we used 80% of all MNChIP-seq peaks as a training subset to assess 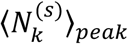 and 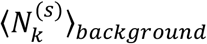. The examples of *k*-mers with the least binding energy calculated using Eq. (1) for two TFs, c-Myc and Otx2, are shown in Figure 2A. As expected, their known binding motifs (8) have the lowest binding energy (Figure 2A).

**Figure 2.**
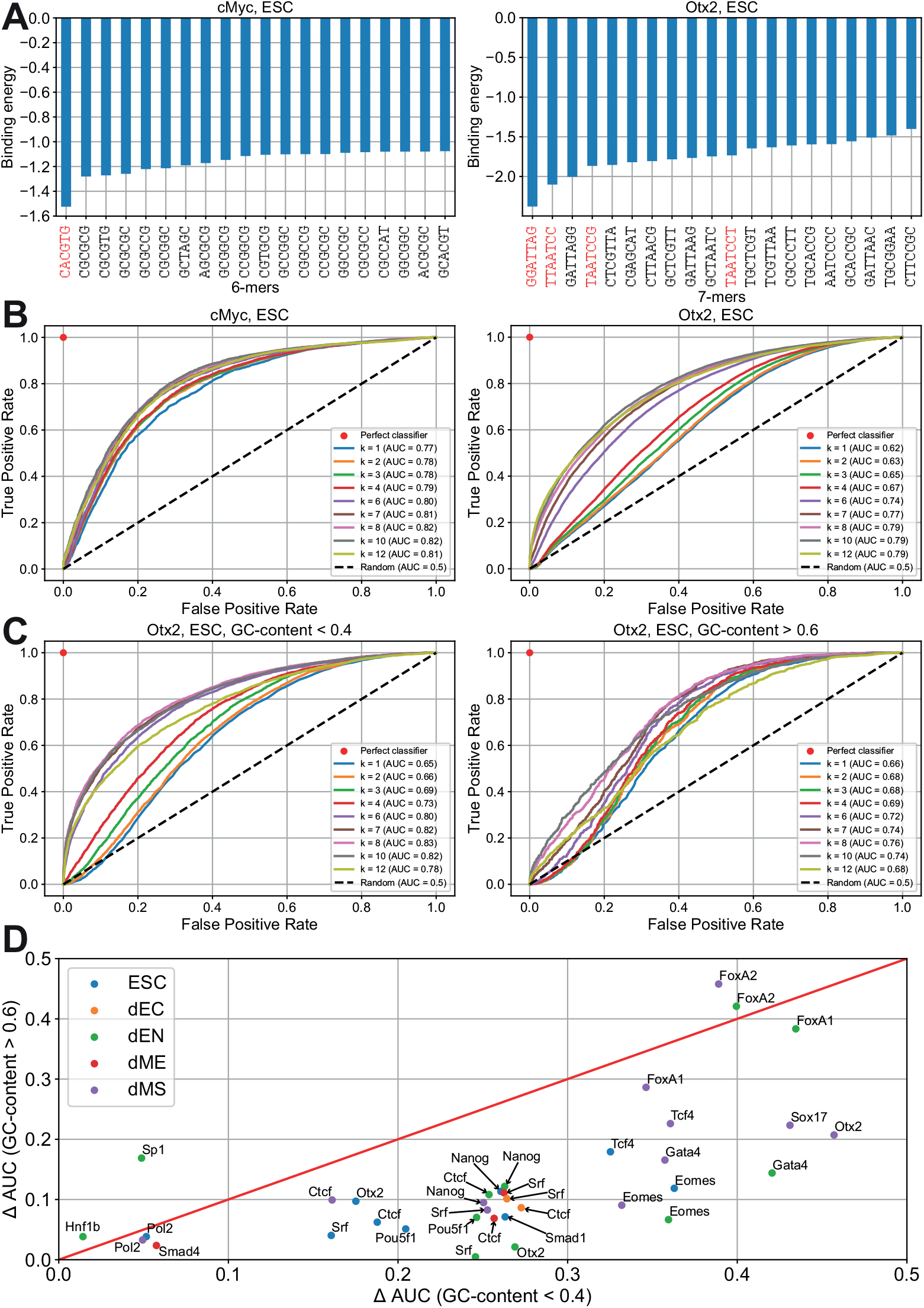
Characterizing *k*-mer specificity of TFs. (**A**) Examples of *k*-mers sorted by increasing binding energy (in the units of *k*_B_*T*) within the proposed peak-over-background energy model for c-Myc (*k*=6) and Otx2 (*k*=7). According to the model, the binding energy of each *k*-mer is defined by Eq. (1). MNChIP-seq data derived from (22), we used 80% of it as a “training subset” to assess binding energies of *k*-mers. The c-Myc binding consensus binding motif, *CACGTG*, is highlighted in red on the left plot, and the 7-mers containing the Otx2 consensus binding motif, *GGATTA*, or its reverse complement, *TAATCC*, are highlighted in red on the right plot. (**B**) Receiver Operating Characteristic (ROC) curves visualizing the proposed energy model’s ability to predict c-Myc and Otx2 binding peaks depending on *k*-mer length. The prediction of whether the sequence belongs to the peak or the background is based on the value of its free energy defined by Eq. (2). Precomputations of binding energies were made using 80% of the MNChIP-seq dataset as described in **A**. The remaining 20% of the dataset was used as a “testing subset” to evaluate the model’s performance. The x-axis shows a false positive rate (FPR), and the y-axis shows a true positive rate (TPR). Pairs of TPR and FPR at different thresholds constitute a ROC curve for each *k*-mer length. The diagonal dotted line signifies classifiers randomly assigning peak status at a consistent probability, equating TPR with FPR. A theoretical perfect classifier is represented by a red dot at the coordinates (0,1). Each ROC curve’s area under the curve (AUC) provides a probabilistic estimate of a randomly selected peak sequence having lower free energy than a randomly selected background sequence, with higher values indicating superior predictive capability. (**C**) Same as B, but instead of analyzing MNChIP-seq peaks of 2 different TFs, we picked out two groups of Otx2 peaks, the first group consisting of peaks with GC-content below 0.4 and the second one consisting of peaks with GC-content above 0.6. All the procedures including splitting into training and testing subsets, *k*-mer binding energy assessment, free energy calculation, and ROC curves visualization were performed separately for each group. (**D**) Summary of *k*-mer binding specificity depending on the GC-content. Using the same procedure as in **C** for each TF and each cell type, we took two groups of MNChIP-seq peaks with GC-content below 0.4 and above 0.6, calculated the area under the ROC curve (AUC) in both groups for different *k*-mer length *k*, and then computed *k*-mer specificity characteristic, ΔAUC, using Eq. (3). The pairs of ΔAUC values for different GC-content were used as coordinates to build a scatter plot. Each point is labeled by the corresponding TF, and the developmental stage is encoded by color. If the number of peaks in either of the two groups is less than 1000, the point is not shown.

The remaining 20% of MNChIP-seq peaks were used as a testing subset to evaluate the model’s performance. To do so, we predicted whether a sequence from the testing subset belongs to the peak or the background based on the value of its free energy (in the units of *k*_B_*T*) defined as:

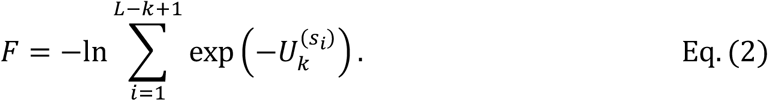

Here, *L* is the length of each sequence and is equal to 100-bp. 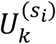 is the precomputed binding energy of a given *k*-mer, *s*_4_, made up from the sequence’s nucleotides with indices from *i* to *i*+*k*-1. The lower the free energy of a sequence is, the more likely the sequence belongs to a binding peak. Notably, in the case of *k*=1, binding energies are assigned to single nucleotides, meaning that the free energy depends on the local GC-content only.

In order to evaluate the predictive ability of the free energy model, Eq. (2), we employ Area under the Curve of the Receiver Operating Characteristic (ROC AUC) (21). Specifically, for any free energy threshold *F*_*t*_ which separates peaks from the background, there is a true positive rate (TPR) which is the fraction of peak sequences having the free energy below this threshold and thus correctly classified as peaks (true positives), and a false positive rate (FPR) which is the proportion of background sequences having the free energy below the threshold and thus misidentified as peaks (false positives). Pairs of FPR and TPR at different thresholds constitute a ROC curve for each length of *k*-mers. The area under the ROC curve (AUC) provides a probabilistic estimate of a randomly selected peak sequence having lower free energy than a randomly selected background sequence. Higher values of AUC indicate a superior predictive capability.

We compared how ROC AUC depends on the *k*-mer length in the case of different TFs (Figure 2B). For TFs with high average GC-content of their binding peaks, such as c-Myc (Figure 1B), there was only a slight increase in ROC AUC computed for *k* > 1 compared to *k* = 1. Therefore, c-Myc represents the case where GC-content provides a dominant contribution to the TF-DNA binding free energy. In contrast, TFs with low average GC-content of their binding peaks, such as Otx2 in ESCs, are characterized by a much larger difference of ROC AUC for *k* > 1 compared to *k* = 1 (Figure 2B).

In the next step, we dissected all Otx2 peaks into two groups. These groups correspond to binding peaks of Otx2 in ESCs with GC-content below 0.4 and above 0.6, respectively. Next, we used these groups as two separate datasets, and evaluated ROC AUC using these datasets (Figure 2C). Here we observed that GC-poor peaks of Otx2 are characterized by a larger difference of ROC AUC between *k* > 1 and *k* = 1 (thus being more *k*-mer-specific), as compared to the case of GC-rich peaks characterized by a smaller difference of ROC AUC between *k* > 1 and *k* = 1 (Figure 2C).

To assess the generality of this phenomenon, we repeated the same procedure with different TFs in different cell types, provided there were at least 1000 peaks in both GC-poor and GC-rich groups. To characterize TF’s *k*-mer specificity with a single value, we introduced the ΔAUC value, defined as the difference between maximal AUC among all *k*-mer lengths and AUC for *k*=1:

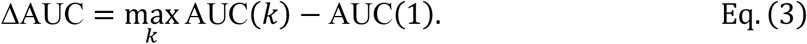

The subtraction of AUC(1) serves the purpose of the GC-content normalization. For each TF and each developmental stage, we calculated ΔAUC using subsets with GC-content below 0.4 and above 0.6, respectively (Figure 2D). We reveal here that the majority of TFs are characterized by larger ΔAUC for GC-poor peaks as compared to GC-rich peaks, suggesting the generality of the discovered phenomenon. Notably, Sp1 and FoxA2 represent two outliers showing slightly higher *k*-mer specificity in GC-rich regions as compared to GC-poor regions (Figure 2D). We emphasize that c-Myc is not represented in Figure 2D due to the fact that the GC-content distribution of c-Myc is shifted towards high GC-content values in all developmental stages (i.e., very few peaks with low GC-content, Figure 1B) and c-Myc is thus characterized by a low *k*-mer specificity (Figure 2B).

### Effect of short tandem repeats on TF-DNA binding specificity

The third local genomic sequence feature that we analyze is short tandem repeats (STRs). In order to test the effect of STRs on TF-DNA binding specificity, we use pair correlation functions developed previously (23-25). These correlation functions represent the probability of finding two nucleotides of a given type separated by relative distance *x*. In particular, for each DNA sequence in a sequence set we compute:

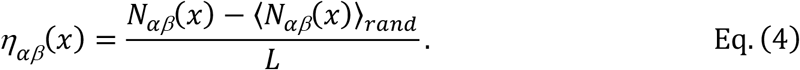

Here, *N*_*αβ*_(*x*) is the total number of nucleotide pairs of the types *α* and *Δ*, separated by the relative distance *x*, ⟨*N*_*αβ*_(*x*)⟩_*rand*_, is the corresponding average number in the randomized sequence set, and *L* is the length of the DNA sequence. The randomization procedure randomly reshuffles each DNA sequence, keeping the GC-content of each sequence intact. The averaging, ⟨*N*_*αβ*_(*x*)⟩_*rand*_, is performed over 100 random realizations of the original sequence. This randomization procedure normalizes the varying genomic GC-content, allowing us to compare symmetry properties of DNA sequences from different genomic locations characterized by a variable average GC-content. At the end of this procedure, for each value of the relative distance *x*, we average the computed *ηαβ*(*x*) over all the DNA sequences in the set.

Here, we analyzed repeat symmetries inside 100-bp-long genomic regions in the center of MNChIP-seq binding peaks of different TFs in different developmental stages, dividing peaks into GC-poor and GC-rich groups, similar to what we performed for the *k*-mer specificity analysis (Figure 3A and B). Notably, the correlation function *η*_CC_(*x*) at *x*=6 (this corresponds to repetitive pattern [CNNNNNC]) provides the strongest distinction between GC-rich and GC-poor regions for a significant number of TFs (Figure 3B). For GC-poor regions, the correlation function *η*_AA_(*x*) is strongly increased at *x*=1 (i.e., enriched in poly(A) tracks) (Figure 3A). In addition, we aligned 2000 bp-long genomic regions near TSSs dividing them into two groups, bound and unbound by the TF, similar to what we performed for the GC-content specificity analysis, and calculated pair correlation functions using 50-bp sliding window (Figure 3C). For the two shown examples of c-Myc and Nanog, *η*_CC_ (*6*) shows an excellent correlation with the mean binding intensity of bound peaks (Figure 3C).

**Figure 3.**
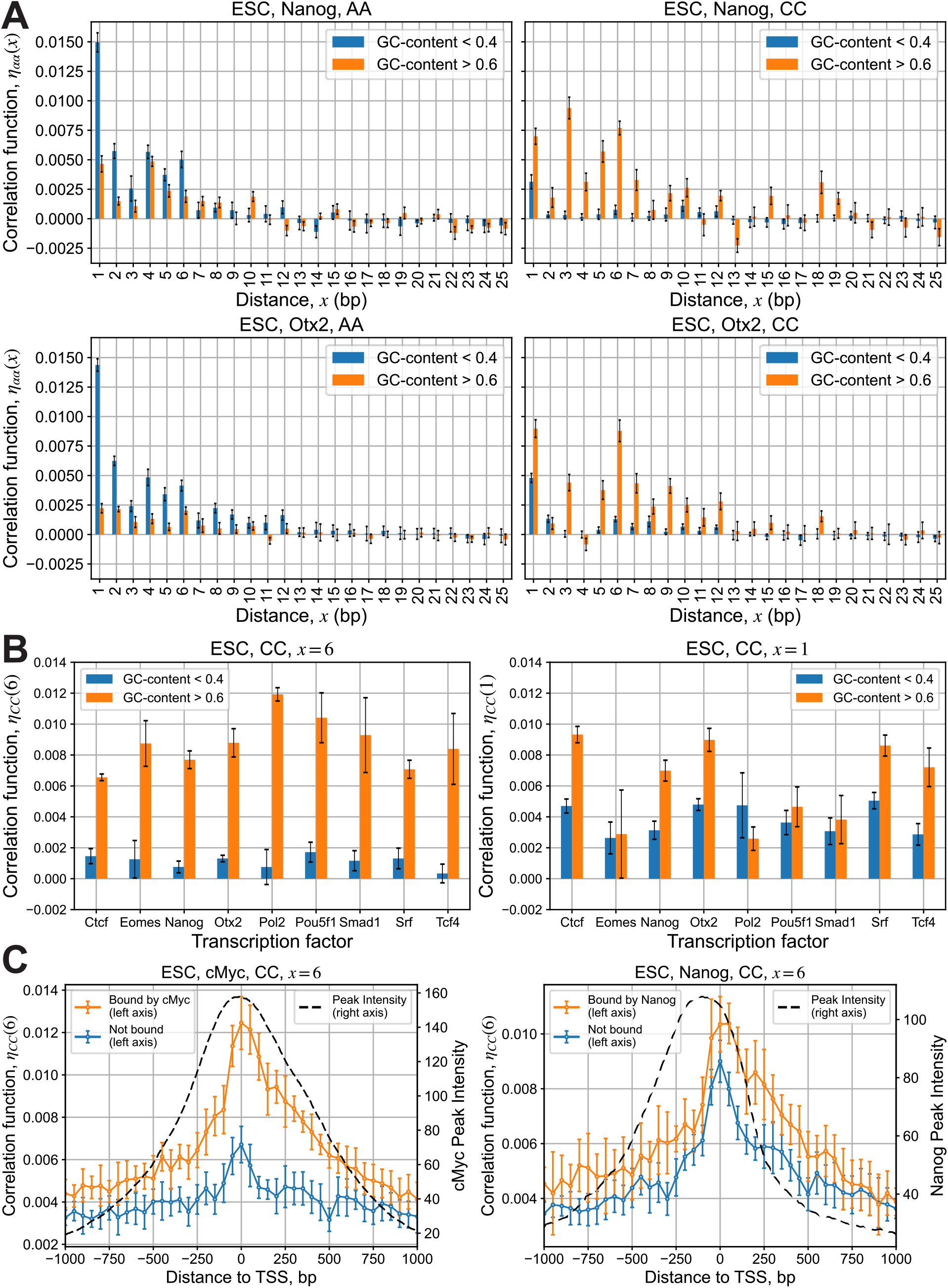
Analysis of genomic repetitive DNA sequence elements depending on the GC-content. (**A**) Correlation functions are computed according Eq. (4) for the two groups of genomic DNA sequences as a function of the relative distance *x* between the pair of nucleotides of a given type (A or C). The first group contains the 100-bp-long genomic sequences of undifferentiated ESCs extracted from the center of MNChIP-seq peaks for Nanog (top) and Otx2 (bottom) with GC-content below 0.4, and the second group contains MNChIP-seq peaks with GC-content above 0.6. To compute error bars for each sequence group, we divide the group into ten subgroups with an equal number of sequences in each subgroup and compute the mean correlation functions, *η*_AA_(*x*) and *η*_CC_(*x*), for each subgroup. Next, we compute the standard deviation of the mean correlation functions between subgroups. Error bars represent one standard deviation in each direction around the mean (i.e., two standard deviations overall). (**B**) Values of the correlation function *η*_CC_(*x*) computed for the same groups as in **A** for various TFs at the selected distances, *x*=6 and *x*=1. The error bars are computed the same way as in **A**. (**C**) The correlation between the mean value of the correlation function, *η*_CC_(6), and the mean binding peak intensity for c-Myc and Nanog near transcription start sites (TSSs). We categorized regions into those bound (orange line) or unbound (blue line) by the TF, defining ‘bound’ as having at least one binding peak within 1000 bp of the TSS, measured from the peak’s nearest edge. The value of *η*_CC_(6) was computed using a 50-bp-wide sliding window and then averaged for regions aligned by the TSS. Dashed line represents the corresponding mean peak intensity for bound TSS regions. Error bars represent one standard deviation of the mean *η*_CC_(6), determined by dividing the genomic regions into 10 equally sized subgroups and calculating the average for each subgroup. For c-Myc, the Pearson linear correlation coefficients between the average peak intensity and *η*_CC_(6): *R* = 0.92 (*p* = 9.1 × 10^−18^) and *R* = 0.82 (*p* = *6*.3 × 10^−11^) for TSS regions bound and unbound by c-Myc, respectively. For Nanog, these values: *R* = 0.*7*1 (*p* = 2.1 × 10^−*7*^) for TSS regions both bound and unbound by Nanog.

Overall, our results demonstrate that for a significant number of TFs, different nonconsensus repetitive DNA sequence elements contribute to TF-DNA binding specificity in GC-rich (the strongest repetitive pattern, [CNNNNNC]/[GNNNNNG]) and GC-poor (the strongest repetitive pattern, poly(A)/poly(T)) genomic regions, respectively.

### Model for competitive Myc-DNA binding with formation of bivalent heterotetramer

Here, we present support for our key working hypothesis that the transitions in the GC-content specificity of key pluripotency factors upon developmental transitions can be explained assuming that they compete with c-Myc for promoter binding, without the need for direct interactions of these TFs with c-Myc. Previously, it has been established that Myc binding to genomic DNA, and its function in transcriptional activation requires heterodimerization with Max (20,26). In a seminal study by Nair and Burley (20), it has been shown that upon DNA binding, c-Myc-Max heterodimer can form a bivalent heterotetramer (Figure 4). AlphaFold3 modeling (19) further supports the formation of a bivalent c-Myc-Max heterotetramer (Figure 4C, Supplementary Table S1). Using c-Myc and Smad1 as a model system, here we show that a simple model for competitive TF-DNA binding qualitatively accounts for the observed distribution of Smad1 between high-GC-content (we term these regions as promoter DNA, DNAP) and low-GC-content (we term these regions as enhancer DNA, DNAE) genomic regions (Figure 5). In particular, in ESCs, the GC-content distribution of Smad1 MNChIP-seq peaks is strongly shifted towards low-GC-content genomic regions (DNAE) (Figure 5A). On the contrary, in dEN the GC-content distribution of Smad1 MNChIP-seq peaks is strongly shifted towards high-GC-content genomic regions (DNAP) (Figure 5B). In sharp contrast with Smad1, the GC-content distribution of Myc peaks is shifted towards high-GC-content genomic regions (DNAP) in both ESCs and dEN (Figure 5A and B). Importantly, while the expression level of Myc is four times higher in ESCs than in dEN, the Smad1 expression level is practically the same in ESCs and dEN (27). Here, we show that the competition between c-Myc bivalent heterotetramer and Smad1 for genomic binding (without assuming any direct interaction between c-Myc and Smad1) can explain the experimentally observed transition of Smad1 genomic preferences upon developmental transition from ESCs to dEN (Figure 5A and B).

**Figure 4.**
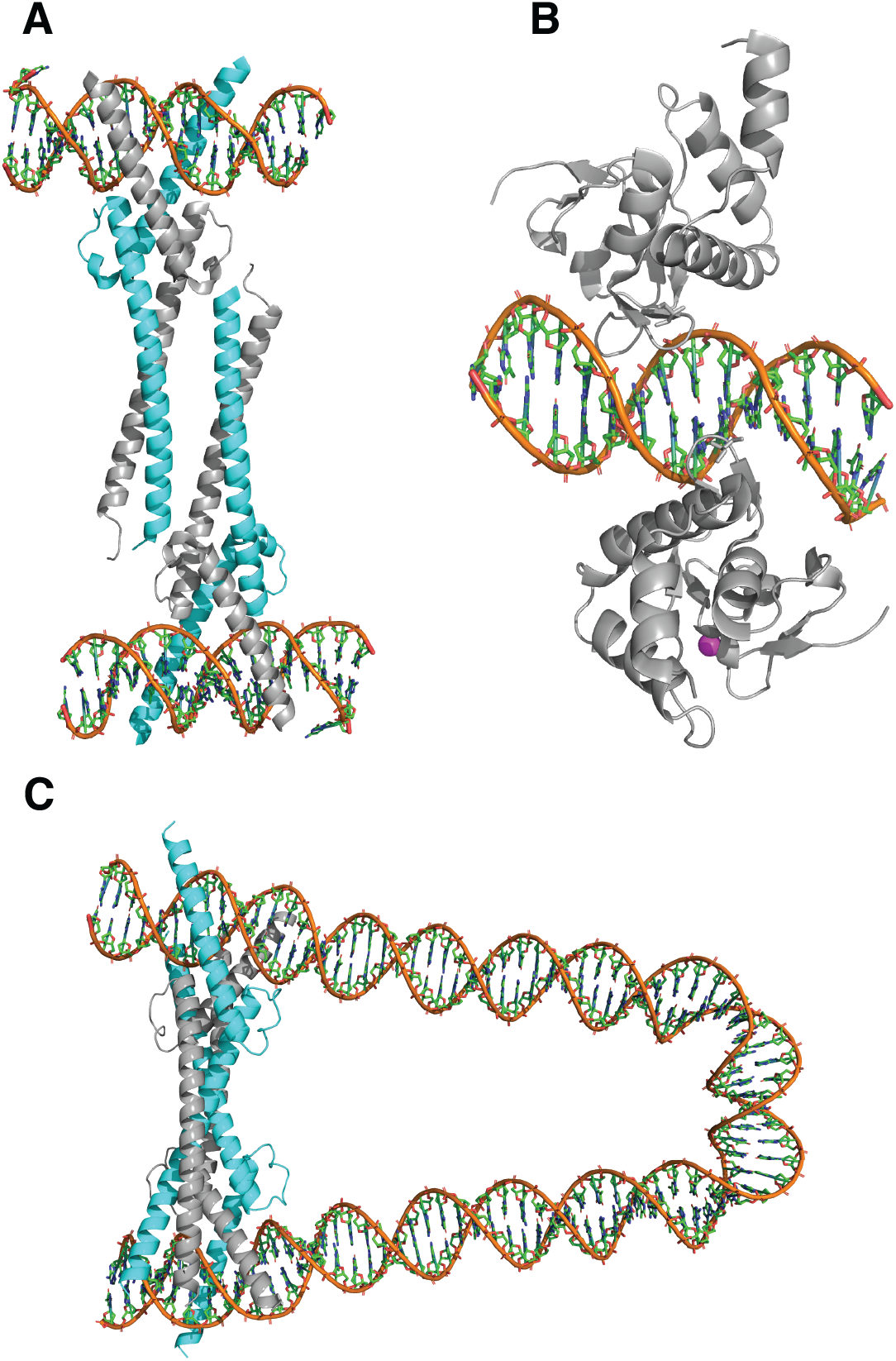
Crystallographic structures of truncated c-Myc-Max bivalent heterotetramer (**A**); and truncated Smad1 homodimer (**B**). The cyan and grey colors in (**A**) represent Myc and Max, respectively. Zink ion is shown in magenta in (**B**). These structures correspond to Protein Data Bank (PDB) accession numbers 1NKP (20) and 3KMP (28), respectively. (**C**) AlphaFold3 prediction of the c-Myc-Max bivalent heterotetramer bound to DNA (sequence ID 5186, Supplementary Table S1), with the length of 96-bp and the GC-content of 54.17%.

**Figure 5.**
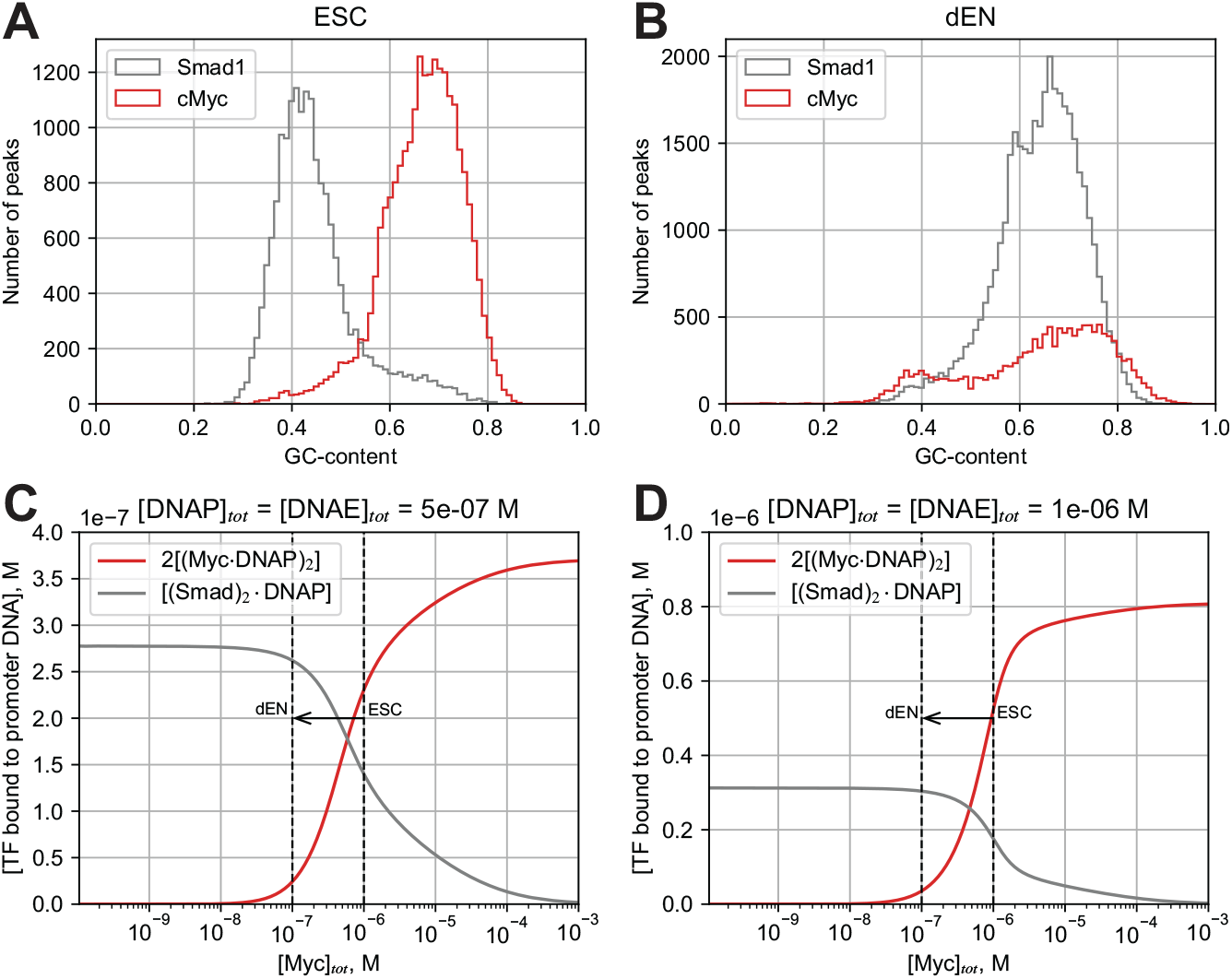
Model for competitive binding of Myc and Smad1 to promoter DNA sequences. (**A, B**) Comparison of GC-content distribution across binding peaks of Myc (red) and Smad1 (grey) in ESCs (**A**) and dEN (**B**). We binned the GC-content range (0 to 1) into 100 equal segments, tallying the occurrence of binding peaks within these intervals. (**C, D**) The computed concentration of the promoter DNA (DNAP) bound by bivalent Myc-Max heterotetramer and by Smad1 homodimer are shown as a function of the total Myc concentration, using two values of the total DNAP and DNAE concentrations, [DNAP]_*tot*_=[DNAE]_*tot*_=0.5 μM (**C**), and [DNAP]_*tot*_=[DNAE]_*tot*_=1 μM (**D**). The following values were used for the remaining model parameters: [Smad]_*tot*_ = 1μM, *K*_1_ = 145nM, *K*_2_ = 90nM, *K*_*E*_ = 500nM, 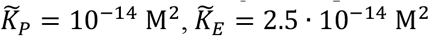 Dashed lines qualitatively represent Myc concentrations in ESCs (right) and dEN (left), respectively. The arrow illustrates the developmental transition from ESCs (with a higher Myc concentration) to dEN (with a lower Myc concentration).

Consider the binding of c-Myc-Max heterodimer to enhancer and promoter DNA sequences. In what follows, we denote c-Myc-Max heterodimer as Myc. The following reactions describe the binding of Myc to DNA:

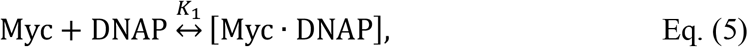

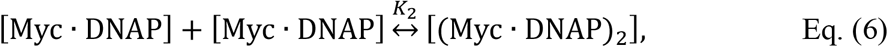

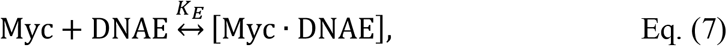

where *K1, K*_2_ and *K*_*E*_ are the equilibrium dissociation constants. Here, [DNAP] is the concentration of the free (i.e., unbound) promoter DNA, [DNAE] is the concentration of the free enhancer DNA, [Myc×DNAP] and [Myc×DNAE] is the concentration of promoter and enhancer DNA bound by Myc heterodimer, respectively; [(Myc×DNAP)_2_] is the concentration of a bivalent Myc heterotetramer (Figure 4). We assume that bivalent Myc heterotetramer can be formed only on DNAP, but not on DNAE. In contrast, we assume that Myc heterodimer can bind both DNAP and DNAE. This constitutes the key assumption of our model.

We now describe competitive Smad1 binding to DNAP and DNAE genomic regions, respectively:

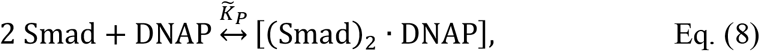

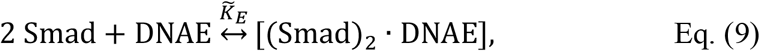

where 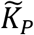 and 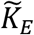 are the equilibrium dissociation constants for Smad1 binding to DNAP and DNAE genomic regions, respectively, and we assume that Smad1 binds both DNAP and DNAE as a homodimer (28).

The corresponding rate equations for Myc:

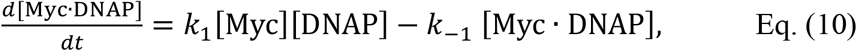

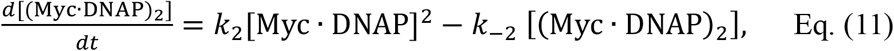

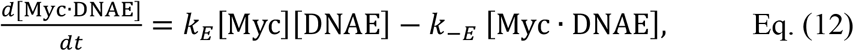

For Smad1 binding we obtain the following:

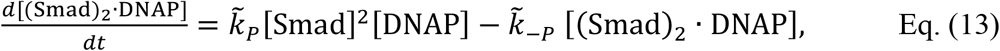

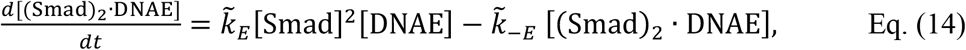

In equilibrium, we obtain:

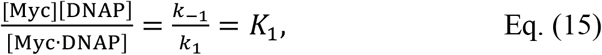

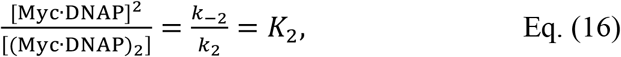

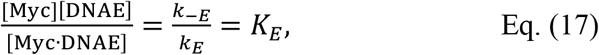

where, *K*_*1*_, *K*_2_ and *K*_*E*_are the equilibrium dissociation constants.

Using Eq. (15) and Eq. (16), we obtain:

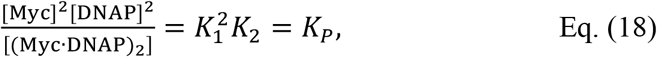

In equilibrium, for Smad1 we obtain the following:

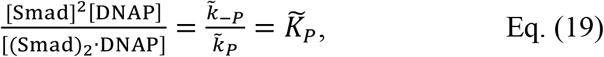

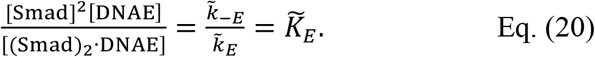

The conservation of mass equations:

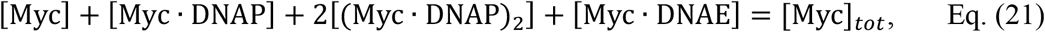

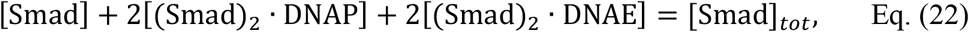

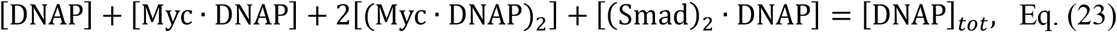

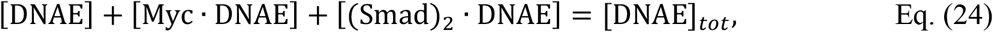

where [Myc]_*tot*_, [Smad]_*tot*_, [DNAP]_*tot*_, and [DNAE]_*tot*_ are the total concentration of Myc, Smad1, DNAP, and DNAE, respectively.

For the equilibrium dissociation constants of Myc, we adopt the following values (20,29): *K*_1_ = 145 nM, *K*_2_ = 90 nM. We assume here for simplicity that all binding sites within DNAP genomic regions can be characterized by the same effective equilibrium constant. Based on the experimental observation that the vast majority of Myc binding events occur in DNAP regions, Figure 5A and B, we assume a weaker Myc binding affinity for DNAE regions as compared to DNAP regions (such weaker binding is characterized by a larger equilibrium dissociation constant), *K*_*E*_ = 500 nM. For Smad1 homodimer binding to DNAP and DNAE we assume 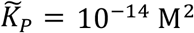 and 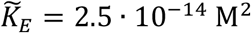, respectively.

Solving the rate equations at equilibrium, we obtain the dependence of the concentration of bivalent Myc heterotetramer and Smad1 homodimer bound to DNAP, as a function of the total Myc concentration, Figure 5C and D. Taking into account that the expression level of Myc in ESCs is four times as high as in dEN, while the expression level of Smad1 is maintained constant in both ESCs and dEN (27), the developmental transition from ESCs to dEN corresponds to the shift from right to left along the *x*-axis in Figure 5C and D. The key observation here is the existence of a sharp transition in the concentration dependence of bound Myc and Smad1, Figure 5C and D. This sharp transition qualitatively reproduces the shift in the GC-content distribution across binding peaks of Myc and Smad1 upon the developmental transition from ESCs to dEN, Figure 5A and B. Our model thus predicts that this transition is solely regulated by the concentration of Myc, competing with Smad1 for promoter binding. The sharpness of the predicted transition stems from the cooperative nature of Myc bivalent heterotetramer binding to promoter regions. Taking together these findings, we predict that first, nonconsensus Myc-DNA and Smad1-DNA binding, and second, the formation of a bivalent Myc heterotetramer on high-GC-content genomic regions, are the two necessary features driving the observed transitions in TF-DNA binding landscape upon developmental transition from ESCs to dEN. It is important to stress that the key prediction of our model is that no direct interaction between Myc and Smad1 is required in order to achieve the transition in the TF-DNA binding landscape. This is consistent with previous findings in seminal work by Orkin et al., revealing that Myc-centered network is largely independent of the core ESC pluripotency network (12). Smad1 belongs to this pluripotency network along with Pou5f1 (Oct4), Nanog, Sox2, Srf, and other TFs (12).

We emphasize that, in contrast to crystallographic measurements (20), AlphaFold3 modeling of a bivalent c-Myc-Max heterotetramer was performed using longer, genomic DNA sequences (Figure 4C, Supplementary Table S1). In this case, c-Myc-Max interaction with DNA leads to DNA looping, resulting in the formation of a bivalent c-Myc-Max heterotetramer (Figure 4C, Supplementary Table S1). In order to validate the generality of this effect, using AlphaFold3, we tested c-Myc-Max binding to 189 genomic sequences with the length varying from 74-bp to 174-bp (Supplementary Table S1). The sequences were selected from c-Myc MNChIP-seq peaks, where each sequence contains two c-Myc specific binding motifs, CACGTG, adjacent to the sequence edges (Supplementary Table S1). We observed that c-Myc-Max forms a bivalent heterotetramer and induces DNA looping for 83% (157 out of 189) of sequences (Figure 4C, Supplementary Table S1). This suggests that a c-Myc-Max bivalent heterotetramer is abundant in the genome.

## Discussion

First, using a statistical-mechanics model for TF-DNA recognition trained on MNChIP-seq data in ESCs, we revealed that key pluripotency TFs possess bimodal intrinsic DNA recognition specificity characterized by fundamentally different mechanisms in low- and high-GC-content genomic regions (Figure 1, Figure 2, and Figure 3). Low-GC-content regions are characterized by enhanced *k*-mer specificity, while high-GC-content regions show enhanced GC-content specificity (Figure 2D). Second, we predicted that c-Myc represents a competitive regulator of TF-DNA binding preferences for key pluripotency factors (Figure 4 and Figure 5). This is consistent with the previous findings revealing that c-Myc-centered network is largely independent of the core ESC pluripotency network containing Smad1, Pou5f1 (Oct4), Nanog, Sox2, Srf, and other TFs (12). In particular, our thermodynamic model predicts that the direct interaction between c-Myc and Smad1 is not essential for the experimentally observed transition in the TF-DNA binding landscape in the course of ESC differentiation (Figure 4 and Figure 5). Rather, the formation of a bivalent c-Myc-Max heterotetramer (for example, by looping DNA) is necessary for achieving this transition (Figure 4C and Figure 5). Past crystallographic studies (20) and our AlphaFold3 modeling provide support for the existence of a bivalent c-Myc-Max heterotetramer (Figure 4). AlphaFold3 prediction of DNA looping by a bivalent c-Myc-Max heterotetramer represents a significant step towards understanding the molecular mechanism of how c-Myc controls gene expression.

Our results point out that c-Myc might affect enhancer-promoter interactions, especially at sub-kilobase length-scales. Strikingly, it was discovered that despite the fact that repression of cohesin eliminates nearly all loop domains, this does not significantly affect the expression level of the majority (∼90%) of genes in a human cell line (30,31). In particular, enhancer-promoter pairs separated by less than 50Kb are weakly affected by a loss of contact domains and loops (31). It was also recently revealed that CTCF and topologically associated domain (TAD) insulation are not required for enhancer-promoter interactions (32). Rather, it was proposed that TF-DNA interactions might be responsible for establishing enhancer-promoter contacts (32). Recently described microcompartments were also shown to be largely unaffected by loss of loop extrusion (33). This suggests that the local, dynamic, three-dimensional conformational ensemble of promoters and enhancer-prompter interactions remains relatively robust despite the elimination of cohesin and CTCF. Our results point out the possibility that transient bivalent (or multivalent) TF-DNA interactions, analogous to a bivalent c-Myc-Max heterotetramer, can stabilize this local, dynamic conformational ensemble of the genome. Consistent with this idea, it was recently identified that p53 drives direct and indirect changes in genome compartments, topologically associated domains (TADs), and DNA loops (34). Further experimental studies probing the effect of how c-Myc affects DNA conformations and enhancer-promoter interactions are necessary in order to validate this hypothesis.

## Supplementary data

Supplementary Data are available Online.

## Acknowledgements

We acknowledge financial support of the Israel Science Foundation (ISF).

