## Supplementary Figures for "Bimodal Specificity of TF-DNA Recognition in Embryonic Stem Cells"

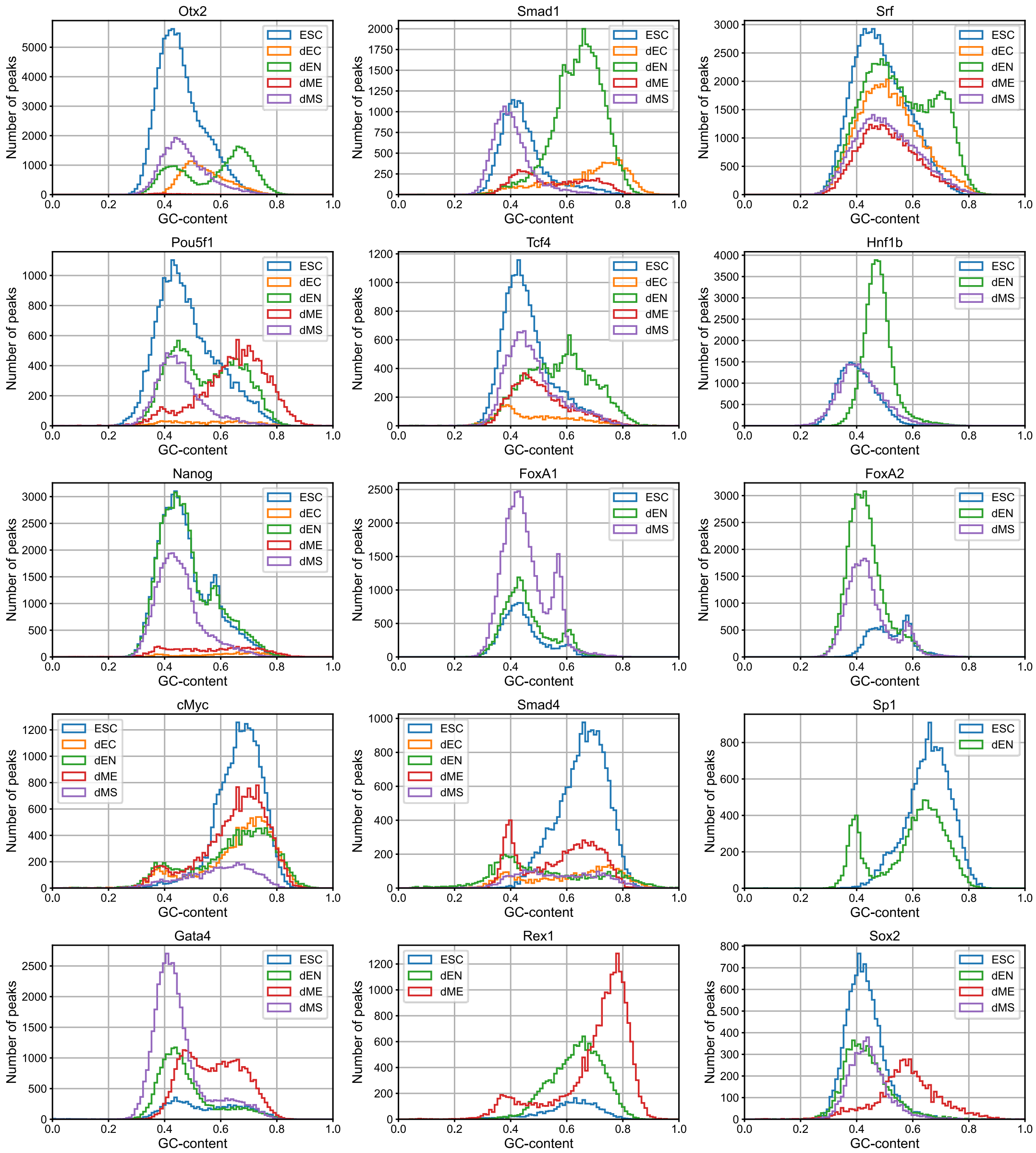


**Supplementary Figure S1**. GC-content distribution of MNChIP-seq peaks for different TFs in five cell types: ESCs, ectoderm (dEC), endoderm (dEN), mesoderm (dME), and mesendoderm (dMS). Each cell type is marked by a unique color. We binned the GC-content range (0 to 1) into 100 equal segments, tallying the occurrence of binding peaks within these intervals. The selected TFs exhibit a bimodality or/and transition of the GC-content distribution from ESCs to other layers. The first two rows include TFs with the shift of the GC-content distribution from lower to higher values upon the developmental transition from ESCs to dEN: Otx2, Smad1, Srf, Pou5f1, Tcf4, and Hnf1b. The third row contains TFs with a similar bimodal GC-content distribution in ESCs and dEN: Nanog, FoxA1, and FoxA2. The fourth row includes TFs with the shift of the GC-content distribution from higher to lower values upon the developmental transition from ESCs to dEN: c-Myc, Smad4, and Sp1. The last row shows TFs with the shift of the GC-content distribution from lower to higher values upon the developmental transition from ESCs to dME: Gata4, Rex1, Sox2.

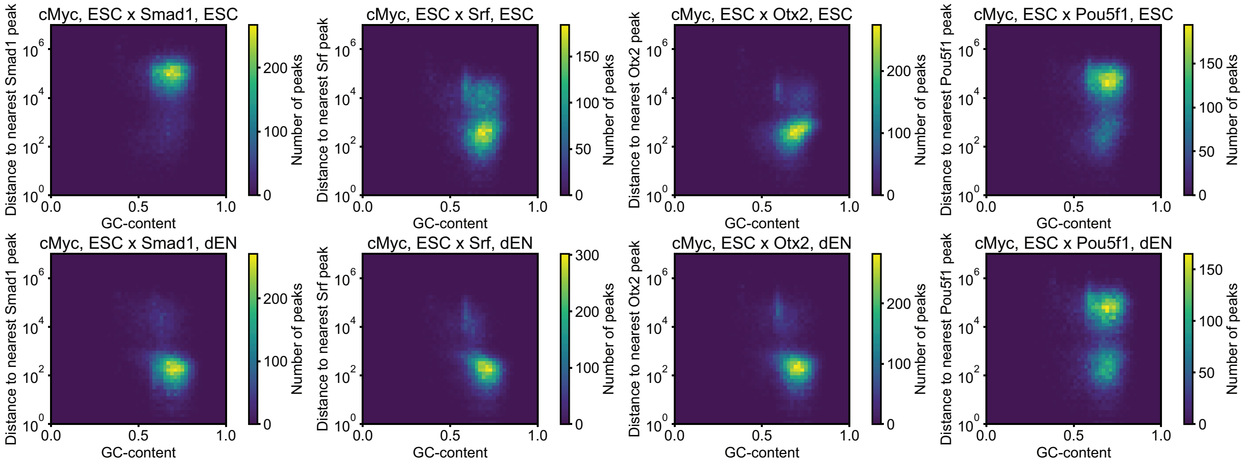


**Supplementary Figure S2**. The joint distribution of the GC-content in c-Myc binding peaks and their proximity to the nearest peaks of Smad1, Srf, Otx2, and Pou5f1, respectively. The upper row shows the joint distribution between peaks of c-Myc in ESCs and peaks of the other TF also in ESCs, and the bottom row shows the joint distribution between peaks of c-Myc in ESCs and peaks of the other TF in dEN. The bottom row demonstrates that Smad1 in dEN occupies genomic regions that were previously bound by c-Myc in ESCs. This means that Smad1 outcompetes c-Myc upon developmental transition from ESCs to dEN. The distances are measured between peak centers. We partitioned the GC-content range (0 to 1) into 50 equal bins and the distance range (1 to $10^7$-bp) into 49 logarithmically scaled bins, counting the frequency of binding peaks within the bins.

### **Supplementary Table** (External Excel File)

**Supplementary Table S1.** (**Sheet 1**) Summary of 189 genomic sequences extracted from c-Myc MNChIP-seq peaks, used in Alphafold3 simulations. The length of the sequences is varying within the interval from 74-bp to 174-bp. Each sequence contains two c-Myc specific binding motifs, CACGTG, adjacent to the sequence edges. The motifs are located 6-bp away from the sequence edges. We observed that c-Myc-Max forms a bivalent heterotetramer inducing DNA looping for 83% (157 out of 189) of sequences. (**Sheet 2**) Sequences used for a control. We randomly reshuffled two sequences (with peak ID 820 and peak ID 1060), producing for each sequence 10 randomized replicas containing the specific binding motif intact, and another 10 randomized replicas reshuffling the entire sequence, including the specific motif. For sequence ID 820, 100% (10 out of 10) randomized replicas containing the intact motif, and 10% (1 out of 10) containing the reshuffled motif formed DNA loops, respectively. For sequence ID 1060, 80% (8 out of 10) randomized replicas containing the intact motif, and 0% (0 out of 10) containing the reshuffled motif formed DNA loops, respectively. (**Sheet 3**) Protein sequences we used in all simulations.
